# A high quality chromosome-level genome assembly for the golden mussel (*Limnoperna fortunei*)

**DOI:** 10.1101/2022.09.29.509984

**Authors:** João Gabriel R. N. Ferreira, Juliana A. Americo, Danielle L. A. S. do Amaral, Fábio Sendim, Yasmin R. da Cunha, The Darwin Tree of Life Project Consortium, Marcela Uliano-Silva, Mauro de F. Rebelo

**Author notes:** Contributed equally to this work (Marcela Uliano-Silva, Mauro de F. Rebelo).

## Abstract

The golden mussel (*Limnoperna fortunei*) is a highly adaptive species that causes environmental and socioeconomic losses in invaded areas. Reference genomes have proven to be a valuable resource for studying the biology of invasive species. While the current golden mussel genome has been useful for identifying new genes, its high fragmentation hinders some applications. In this Data Note, we provide the first chromosome-level reference genome for the golden mussel. The genome was built using Hi-C, PacBio HiFi and 10X sequencing data. The final assembly contains 99.4% of its total length assembled to the 15 chromosomes of the species and a scaffold N50 of 97.05 Mb. Approximately 47% of the genome was annotated as repetitive sequences. A total of 34 862 protein-coding genes were predicted, of which 84.7% were functionally annotated. This new high quality genome is expected to support both basic and applied research on this invasive species.

**Species taxonomy:** Eukaryota; Opisthokonta; Metazoa; Eumetazoa; Bilateria; Protostomia; Spiralia; Lophotrochozoa; Mollusca; Bivalvia; Autobranchia; Pteriomorphia; Mytilida; Mytiloidea; Mytilidae; Arcuatulinae; Limnoperna; Limnoperna fortunei (Dunker, 1857) (NCBI Taxonomy ID: 356393)

## Background

*Limnoperna fortunei* — popularly known as the golden mussel — is a freshwater bivalve species native to Southeast China which has successfully established itself as an invasive species in other Asian countries (Cambodia, Japan, Laos, South Korea, Taiwan, and Thailand) and in several South American countries (Argentina, Brazil, Paraguay, and Uruguay). Because of its impact on ecosystem structure and function, the golden mussel is considered an efficient ecosystem engineer, and its establishment is associated with changes in the proportions of species of the local fauna [1, 2]. Current control strategies have not proven to be effective and the species continues to spread [3].

Reference genomes are an important resource for the study of invasive species. They have been used to study invasion dynamics, identifying molecular mechanisms conferring adaptiveness as well as promising genes for biotechnology-based control strategies [4]. There currently is a genome assembly for the golden mussel [5], but it is a highly fragmented representation of the 15 chromosomes (2n=30) of the species [6, 7] assembled mostly based on Illumina sequencing reads. Limitations of the Illumina-based genome constrain its applications in resequencing and comparative genomic studies.

Recent advances in sequencing technologies and bioinformatics algorithms have made the development of high quality reference genomes scalable and affordable. In this Data Note, we present such a high quality reference genome developed for the golden mussel. This new genome is expected to be a valuable reference for future studies that assess the genetic diversity of the species, postulate genomic evolution by comparison to other bivalve species, and characterize gene families of interest, among other goals.

## Genome assembly report

The size of the final genome assembly is 1.34 Gb; 99.24% of its total length is distributed over the 15 largest scaffolds (Figure 1), which correspond to the haploid chromosome number (n=15) of the species. The largest contig and the largest scaffold are 8.3 Mb and 115 Mb long, respectively. The genome GC content is 33.6% (Supplementary Table S1).

**Figure 1.**
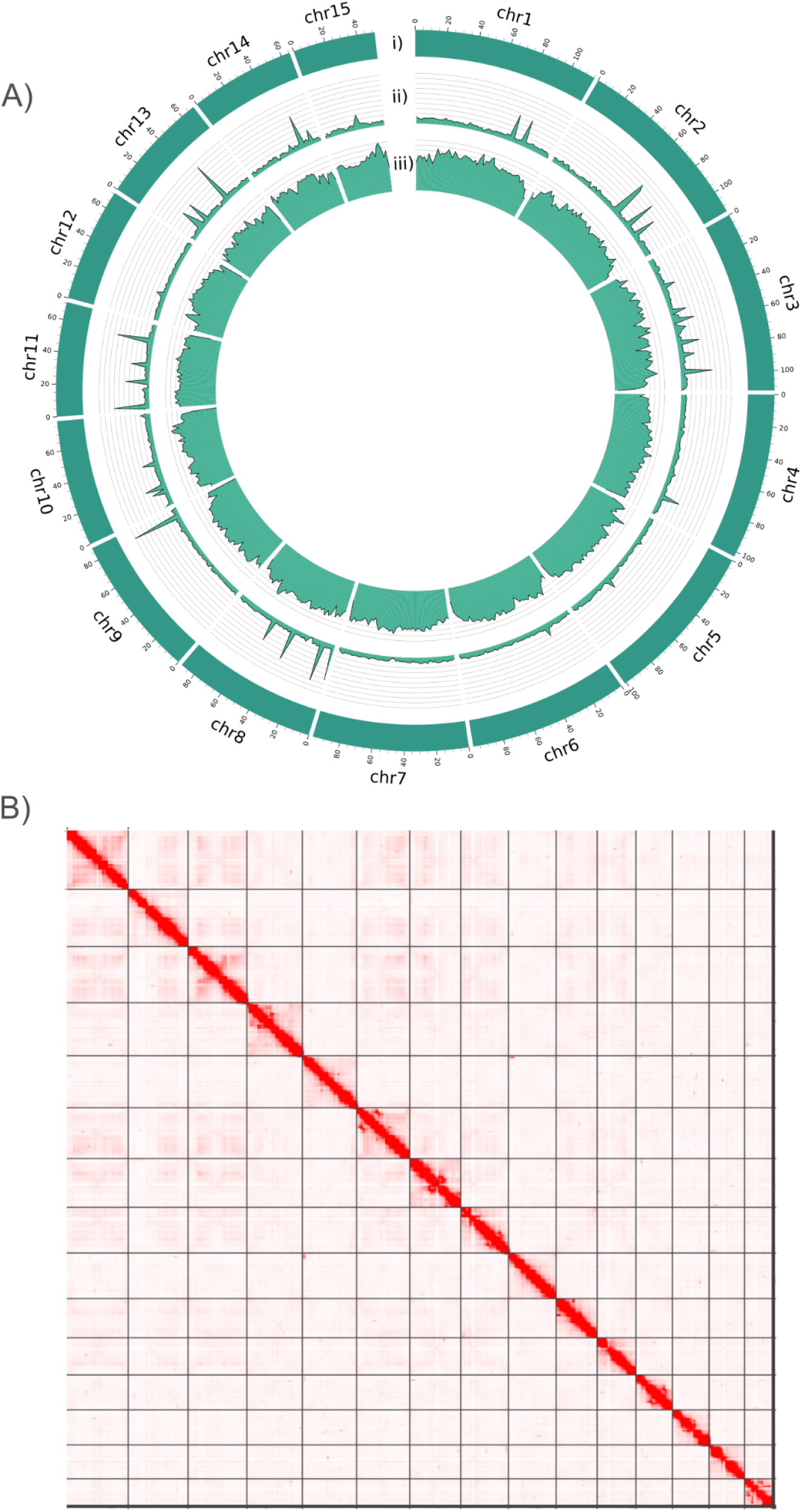
The genome landscape. A) Circos representation of the 15 chromosomes assembled in this study. Each track represents: i) the size of each chromosome, ii) the gene density, and the iii) repeat density over the chromosome sequences, calculated using a 2 Mb window size. B) Hi-C contact map with chromosomes displayed in size order from top to bottom and from left to right.

Table 1 presents genomic statistics of the Illumina-based assembly and the new chromosome-level reference produced in this study. The new reference scaffold N50 is 313-fold greater than its Illumina-based predecessor (Table 1). An improvement has also been achieved in genome completeness as shown by an increase in the percentage of BUSCO genes (Table 1). The QV score of 53 of the chromosome-level genome represents a base call accuracy of 99.999%. All the quality metrics calculated for the new assembly conform to the standards of the Vertebrates Genome Project (VGP) for what is considered a high-quality genome [8].

**Table 1.** Comparison of assembly metrics between the Illumina-based and the new golden mussel genome.

|  | **Illumina-based genome**<br>**(GCA_003130415.1)** | **Chromosome-level genome**<br>**(GCA_944474755.1)** |
| --- | --- | --- |
| **Total assembly length**<br>**(Gb)** | 1.67 | 1.34 |
| **GC content (%)** | 33.6 | 33.8 |
| **Number of scaffolds** | 20 580 | 309 |
| **Scaffold N50 (Mb)** | 0.31 | 97.05 |
| **Scaffold L50** | 1 489 | 7 |
| **Number of contigs** | 61 175 | 1 838 |
| **Contig N50 (Mb)** | 0.03 | 1.50 |
| **Contig L50** | 16 521 | 277 |
| **QV** | 14.89 | 53.36 |
| **BUSCO**<br>**(metazoa_odb10)** | C:66.8% [S:65.0%,D:1.8%],<br>F:19.3%,M:13.9%,n:954 | C:95.6% [S:95.0%,D:0.6%],<br>F:2.2%,M:2.2%,n:954 |
| **BUSCO**<br>**(mollusca_odb10)** | C:56.0% [S:54.7%,D:1.3%],<br>F:7.4%,M:36.6%,n:5295 | C:87.0% [S:86.0%,D:1.0%],<br>F:3.2%,M:9.8%,n:5295 |

BUSCO statistics. C=complete; S=complete and single-copy; D=complete and duplicated; F=fragmented; M=missing; n=number of BUSCO genes from reference dataset.

## Repeat annotation

Almost half (46.93%) of the genome was annotated as repetitive sequences according to the EarlGrey pipeline [9], with 35.80% of the genome labeled as unclassified repeats (Table 2). Similarly high proportions of unclassified repeats have been reported in other mussels [10, 11]. The second most frequent repeat class detected was Long interspersed nuclear elements (LINE), representing 4.51% of the total genome.

**Table 2.** Repetitive elements identified in the golden mussel genome.

| Classification* | Total sequence length (bp) | Sequences count | Proportion of genome (%) | Number of distinct classifications |
| --- | --- | --- | --- | --- |
| DNA | 46 269 487 | 64 938 | 3.46 | 201 |
| LINE | 60 245 208 | 64 949 | 4.51 | 224 |
| LTR | 28 342 781 | 50 395 | 2.12 | 112 |
| Other<br>(Simple Repeat,<br>Microsatellite,<br>RNA) | 216 907 | 244 | 0.02 | 2 |
| Penelope | 11 115 342 | 24 065 | 0.83 | 23 |
| Rolling Circle | 1 640 436 | 1 503 | 0.12 | 6 |
| SINE | 831 095 | 723 | 0.06 | 3 |
| Unclassified | 478 115 783 | 883 466 | 35.80 | 1 494 |

LINE=Long interspersed nuclear elements; LTR=Long terminal repeats; SINE=Short interspersed nuclear element
\* Classification in alphabetical order.

## Gene prediction and functional annotation

A total of 34 862 protein-coding genes were predicted by the Ensembl rapid annotation pipeline [12], with 68 899 proteins inferred. Most genes (53.5%) were associated with a single protein, with about 21.8% associated with two proteins, and 24.7% with three or more proteins (Supplementary Table S2). In addition to the protein-coding genes, 58 911 non-coding genes were predicted, most of which (56.5%) were classified as long non-coding RNA (lncRNA) (Table 3).

**Table 3.** Categories of predicted genes.

| Statistic | Value |
| --- | --- |
| Protein-coding genes | 34 862 |
| Non-coding genes | 58 911 |
| lncRNA | 33 258 |
| Y_RNA | 9 316 |
| tRNA | 7 582 |
| ribozyme | 5 091 |
| misc_RNA | 1 641 |

**Table 3.** (continued).
|  |  |
| --- | --- |
| rRNA | 1 410 |
| snRNA | 565 |
| snoRNA | 47 |
| scaRNA | 1 |

lncRNA=long non-coding RNA; tRNA=transfer RNA; misc\_RNA=miscellaneous RNA; rRNA=ribosomal RNA; snRNA=small nuclear RNA; snoRNA=small nucleolar RNA; scaRNA=small Cajal body-specific RNA.

**Table 4.** Gene prediction statistics.

| Statistic | Value |
| --- | --- |
| Average gene length (bp) | 9 426 |
| Protein-coding genes (bp) | 17 765 |
| Non-coding genes (bp) | 4 492 |
| Exons | 719 821 |
| Average exon length (bp) | 229 |
| Proteins | 68 899 |
| Average protein length (aa) | 462 |
| Gene density (No. genes/100kb) | 7.02 |
| Protein-coding genes | 2.61 |
| Non-coding genes | 4.41 |

Functional annotation was performed against public databases. Of the 34 862 protein-coding genes, 19 899 (57.08%) had at least one hit against the curated SwissProt database (Figure 2). The genes were also searched against Pfam-A for protein domain annotation, with 20 963 (60.13%) genes showing at least one Pfam hit. Finally, genes were mapped against the eggNOG database to retrieve gene ontology (GO) and KEGG pathway annotations. Of the 34 862 genes, 6 183 (17.74%) were associated to a KEGG pathway and 9 746 (27.96%) to at least one GO term.

**Figure 2.**
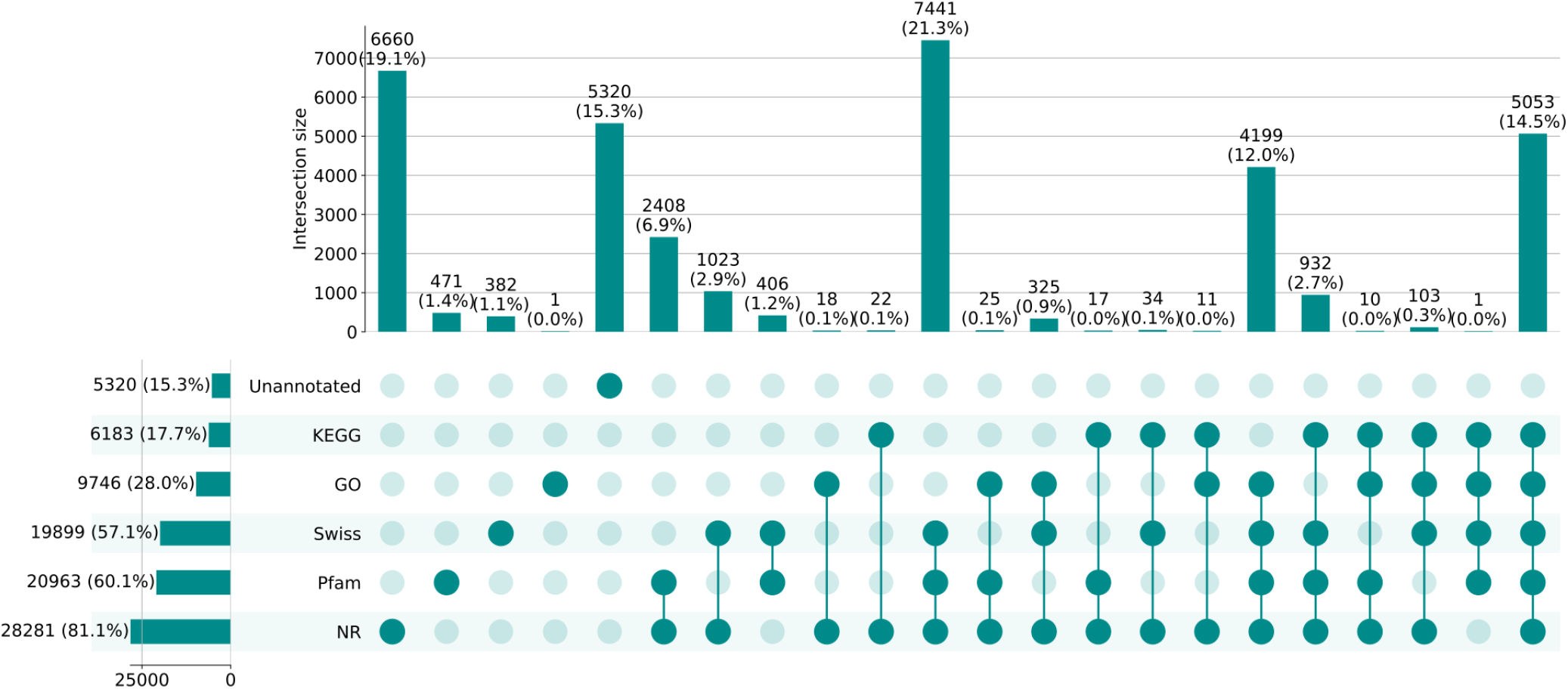
UpSetPlot representing the different functional annotations. Horizontal bars represent the total number of genes annotated according to each database. Vertical bars represent overlapping annotations (i.e. genes annotated by a single or a combination of databases), as indicated by the connected dark green circles.

## Methods

### Sample collection

Golden mussel specimens were collected from Taquari River, São Paulo, Brazil (23°16’45.7”S 49°12’01.7”W) on March 17, 2021. Three representative specimens were deposited in the molluscan collection of the National Museum administered by the Federal University of Rio de Janeiro (identification numbers: IB UFRJ 19950, IB UFRJ 19952 and IB UFRJ 19954). Other specimens were taxonomically identified by Dr. Igor Christo Miyahira. Finally, a set of specimens had their tissues – gonads, adductor muscle, digestive gland, gills, and foot – dissected and preserved in dry ice at -80ºC until and during transportation to the Wellcome Sanger Institute (WSI) in Hinxton, Cambridgeshire, United Kingdom for further processing and sequencing.

### DNA extraction

DNA extraction was performed at the WSI’s Tree of Life laboratory. Golden mussel samples were weighed and disrupted using a Covaris cryoPREP Automated Dry Pulveriser which subjects tissue – gill tissue was selected – to multiple impacts until it becomes a fine powder. Twenty-five mg of this powder was used for DNA extraction and 50 mg was set aside for Hi-C. DNA extraction was performed using a Qiagen MagAttract HMW DNA extraction kit on a KingFisher APEX. Fifty ng of DNA was submitted for 10X sequencing with any low molecular weight DNA removed prior to sequencing using a 0.8X AMpure XP purification kit. Similarly, prior to submission for PacBio sequencing, high molecular weight DNA was sheared to an average fragment size of between 12 kb and 20 kb using a MegaRuptor 3 (speed setting 30). The sheared DNA was purified by solid-phase reversible immobilization using AMpure PB beads with a 1.8X ratio of beads to sample. The concentration of sheared DNA was assessed using a Qubit Fluorometer with Qubit dsDNA High Sensitivity Assay kit and Nanodrop spectrophotometer, while the fragment size distribution was assessed using an Agilent FemtoPulse.

### Sequencing

All sequencing libraries were constructed using DNA extracted from a single specimen, a female golden mussel identified as xbLimFort5. Pacific Biosciences HiFi circular consensus and 10X Genomics linked-reads sequencing libraries were constructed according to the manufacturers’ instructions. Sequencing was performed by the Scientific Operations core at the Wellcome Sanger Institute on Pacific Biosciences SEQUEL II (HiFi) and Illumina NovaSeq (10X) instruments. Hi-C data were generated using the Arima v2.0 kit and sequenced on a NovaSeq 6000 instrument.

### Genome assembly

The genome assembly pipeline is summarized in Figure 3. The initial set of contigs was assembled using HiFiasm v0.16.1 combining HiFi and Hi-C reads in the Hi-C integrated mode [13]. 10X linked-reads were mapped to contigs using LongRanger v2.2.2 [14] and then Freebayes v1.3.1 [15] was used to polish the contigs based on the 10X mapping. The polished contigs were then scaffolded using the YaHS pipeline v1.0 [16]. Finally, scaffolds were manually curated by WSI’s Genome Reference Informatics Team (GRIT) following the protocol described by Kerstin and colleagues (2021) [17]. The curated scaffolds represent the final genome assembly, which was then annotated using Ensembl Rapid Annotation Pipeline [12].

**Figure 3.**
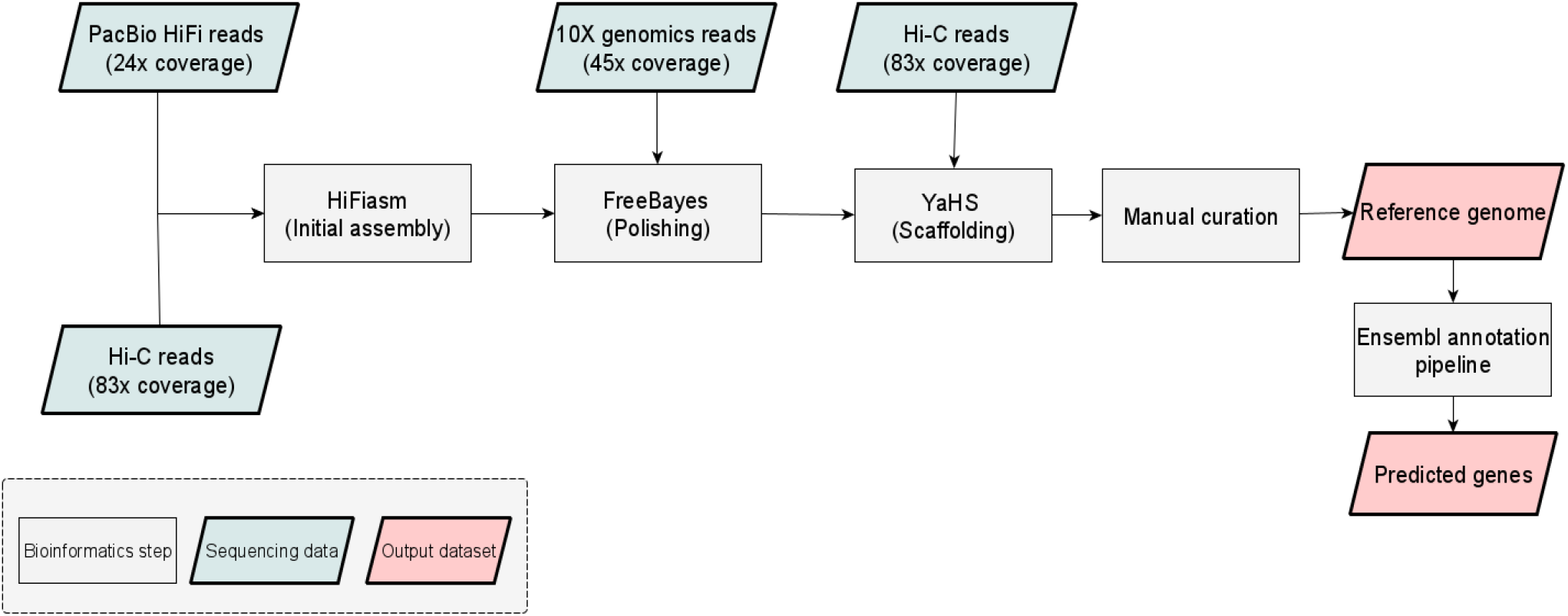
Genome assembly pipeline.

### Genome assembly statistics

General assembly statistics were calculated using an internal perl script. Completeness stats were calculated using BUSCO [18] version 5.0 with two datasets: metazoa_odb10 and mollusca_odb10. QV statistics was calculated using the Merqury software [19] using HiFi data (for the new genome) and Illumina paired-end data (for the Illumina-based genome).

### Functional annotation

The longest protein inferred from each gene was selected using the primary_transcript.py script from OrthoFinder v2.5.4 [20]. Those proteins were aligned against the SwissProt database (downloaded on June 2, 2022) using BLASTP v2.12.0+ from blast+ package [21] and against the NR database (downloaded on June 24, 2022) using Diamond v2.0.15.153 [22]. Both alignments were done using a threshold of 1e^-5^ for the e-value parameter. The eggNOG mapper v2 web server [23] was used to attribute GO terms and KEGG pathways to each protein. Alignment against Pfam was done using the *hmmsearch* (e-value threshold of 1e^-5^) command from HMMER v3.3.1 [24] to annotate protein domains associated with the predicted genes. Sequences were labeled as “unannotated” when they didn’t have a hit to any of the five databases searched (NR, SwissProt, GO, KEGG and Pfam).

## Data availability

The genome sequence is available in the NCBI under accession GCA_944474755.1, while contigs representing the alternative haplotype are available as GCA_944589985.1. Raw data accessions are summarized in Table 5.

**Table 5.** Accession numbers of raw sequencing data used for the genome assembly project.

| Library | Accession(s) |
| --- | --- |
| Pacific Biosciences SEQUEL II (HiFi) | ERR9713989-91, ERR9713993 |
| 10X Genomics Illumina | ERR9503462-65 |
| Hi-C Illumina | ERR9503466 |

## Acknowledgements

This work was financed by the Brazilian National Electric Energy Agency ANEEL R&D program (grant PD-10381-0419/2019). We also thank CTG Brasil, Tijoá Energia and Spic Brasil for funding this project through the ANEEL R&D Program. João Gabriel R. N. Ferreira and Fábio Sendim were recipients of Ph.D fellowships and Yasmin R. da Cunha was a recipient of Master’s fellowship from CAPES, a federal government agency of the Brazilian Ministry of Education which supports graduate students and faculty. Genome sequencing and assembly was provided by the Wellcome Sanger Institute Tree of Life Programme in collaboration with the Bio Bureau Biotechnology company.

## Ethics/compliance issues

The materials that have contributed to this Data Note have been supplied by a Darwin Tree of Life Partner. The submission of materials by a Darwin Tree of Life Partner is subject to the Darwin Tree of Life Project Sampling Code of Practice (https://www.darwintreeoflife.org/wp-content/uploads/2021/03/DToL-Sampling-Code-of-Practice.pdf). By agreeing with and signing up to the Sampling Code of Practice, the Darwin Tree of Life Partner agrees they will meet the legal and ethical requirements and standards set out within this document in respect of all samples acquired for, and supplied to, the Darwin Tree of Life Project. Each transfer of samples is further undertaken according to a Research Collaboration Agreement or Material Transfer Agreement entered into by the Darwin Tree of Life Partner, Genome Research Limited (operating as the Wellcome Sanger Institute), and in some circumstances other Darwin Tree of Life collaborators.

