## Supplementary Data for "A high quality chromosome-level genome assembly for the golden mussel (*Limnoperna fortunei*)"

**Supplementary Table S1**. General statistics for the golden mussel genome sequence.

COMPOSITION A = 442398358 (33.1%), C = 224258371 (16.8%), G = 224472855 (16.8%), T = 444124758 (33.3%), N = 305617 (0.0%), CpG = 52146080 (3.9%)

SCAFFOLD sum = 1335559959, n = 309, mean = 4322200.51456311, largest = 115440917, smallest = 1000
SCAFFOLD N50 = 97051362, L50 = 7
SCAFFOLD N60 = 89737950, L60 = 8
SCAFFOLD N70 = 78518013, L70 = 10
SCAFFOLD N80 = 70815894, L80 = 11
SCAFFOLD N90 = 69091614, L90 = 13
SCAFFOLD N100 = 1000, L100 = 309

CONTIG sum = 1335254342, n = 1838, mean = 726471.350380849, largest = 8 315 943, smallest = 273
CONTIG N50 = 1498882, L50 = 277
CONTIG N60 = 1221040, L60 = 376
CONTIG N70 = 962899, L70 = 500
CONTIG N80 = 718770, L80 = 662
CONTIG N90 = 441659, L90 = 895
CONTIG N100 = 273, L100 = 1838
GAP sum = 305617, n = 1529, mean = 199.880313930674, largest = 200, smallest = 17

**Supplementary Table S2**. Number of proteins associated with each gene.

| Number of proteins per gene | Number of genes |
|---|---|
| 1 | 18 647 |
| 2 | 7 597 |
| 3 | 4 150 |
| 4 | 2 210 |
| 5 | 1 124 |
| 6 | 535 |
| 7 | 281 |
| 8 | 131 |
| 9 | 89 |
| 10 | 31 |
| 11 | 36 |
| 12 | 15 |
| 13 | 7 |
| 14 | 4 |
| 15 | 2 |
| 19 | 2 |
| 21 | 1 |
